# The velocity-curvature power law in *Drosophila* larval locomotion

**DOI:** 10.1101/062166

**Authors:** Myrka Zago, Francesco Lacquaniti, Alex Gomez-Marin

**Author notes:** **Summary statement.** The motor control power law relating speed and curvature in humans is at work in the humble maggot.

## Abstract

We report the discovery that the locomotor trajectories generated by crawling fruit fly larvae follow the same power law relationship between speed and curvature previously found in the human motor control of hand-drawing, walking, eye movements and speech. Using high resolution behavioral tracking of individual flies in different sensory environments, we tested the power law by making maggots trace different trajectory types in naturalistic conditions, from reaching-like movements to scribbles. In all these conditions, we found that the law holds, and also that the exponent of the larval scaling law approaches 3/4, rather than the usual 2/3 exponent found in almost all human situations. This is consistent with recent findings on humans drawing ellipses on water, where dynamic effects related to medium viscosity have been shown to increase the exponent that would emerge from purely kinematic-geometric constraints. To our knowledge, the speed-curvature power law has only been studied in human and non-human primates, our work then being the first demonstration of the speed-curvature scaling principle in other species. As there are still different competing hypotheses for the origin of such law in humans (one invoking complex cortical computations in primates; another postulating its emergence from the coupling of viscoelastic muscle properties with simple central pattern generation) our findings in the larva demonstrate that the law is possible in an animal with a nervous system orders of magnitude simpler than that of humans, thus supporting the latter view. Given that our discovery is in *Drosophila* (amenable to precise genetic manipulations, electron microscopy reconstruction of neural circuits, imaging in behaving animals, electrophysiology, and other techniques) this opens great potential for uncovering the mechanistic implementation of the velocity-curvature power law. Such scaling laws might exist because natural selection favors processes that remain behaviorally efficient across a wide range of contexts in distantly related species. Our work is an effort to search for shared principles of animal behavior across phyla.

## Introduction

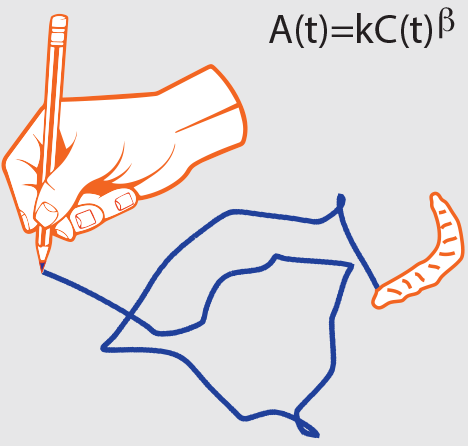

When we scribble our name on a piece of paper, the instantaneous angular velocity is related to the local path curvature following a power law [1]. Being one of the best-studied characteristics of human voluntary movements, the law was originally described for hand-drawing movements across a wide range of sizes, speeds, and curvatures [1–2],and subsequently has been observed also in smooth eye movements, speech, and walking [3]. Movements complying with the law are optimal, with maximum smoothness and minimum variability [2–3]. The law is not dictated by physical constraints: even when the path is imposed, as in hand-drawing, movement speed could in principle vary in infinite ways. Therefore, the law must result from physiological constraints, although its origin remains debated. According to one view, the law originates by decoding complex neural processes at cortical level; indeed, population vectors in motor cortex obey the power law during drawing [4]. According to another view, the law stems from simple harmonic motions—such as those output by spinal Central Pattern Generators— coupled with the viscoelastic properties of muscles [5]. To our knowledge, the power law has only been studied in human [1–3] and non-human[4] primates.

Here we report that *Drosophila* melanogaster larvae, whose movements are controlled by a much simpler neural system [6], display a velocity-curvature power law while crawling. This demonstrates that the law can emerge from the interplay between simple neural commands and biomechanics. Our findings support the view that, despite huge divergence in anatomy, functional complexity and ecological contingencies, basic principles of motor control resulting in efficient behavior are shared across distantly related species [3,7].

## Materials and Methods

**Experimental Procedures.** Experimental procedures and behavioral tracking of *Drosophila* larva behavior were the same as those described in [8]. Third-instar *Drosophila* melanogaster larvae inthe foraging stage were washed in a 15%-sucrose solution and then transferred to a rectangular, flat-lid arena coated with a 3%-agarose slab. Lids, agarose layers and individual larvae were not reused. Animals were tracked at a resolution of 7 frames per second at 90 μm per pixel for five minutes. Tracking was interrupted if the animal touched the plate boundaries. Custom-made, freely-available tracking scripts [9] extracted the location of the centroid of the animal (as well as other points of interest such as the head or tail) from postural sequences. As explained in the main text, we used 3 groups of larvae exposed to different experimental conditions: overshoot, approach,and dispersal. Odor gradients were created by suspending a 10 μl droplet of ethyl-butyrate from the top of the arena at a position out of reach of the animal.

**Main data analysis.** The raw x, y position samples of the centroid (output by the tracking software, see above) were low-pass filtered (second-order, zero-phase-lag Butterworth filter) with a cut-off frequency of 0.07 Hz. This cut-off frequency was chosen after computing the power-spectrum density of the raw (unfiltered) data and verifying that the power was approximately flat up to about 0.01 Hz, and then dropped rapidly. At 0.07 Hz, the power was always down by about 30 dB. We then interpolated the filtered data with cubic splines, computed the time derivatives of the interpolating spline, the curvature C(t) and angular velocity A(t) from standard differential geometry [10].Least-squares orthogonal regression of log A(t) versus Log C(t) was performed to estimate the exponent β and the variance accounted for r2 by the power law A(t)=kC(t)^β^. Statistically significant differences of the exponent β between experimental conditions were assessed using non-parametric tests (Kruskal-Wallis ANOVA by ranks followed by multiple comparisons), because the data were not normally distributed (Kolmogorov-Smirnov test).

**Additional analyses and results.** To assess the effects of low-pass filtering the data [11], we performed the orthogonal regression of log A(t) versus Log C(t) over a wide range of frequency cutoffs, including no filtering whatsoever. **Figure S1** reports the results for the overshoot condition, but very similar results were obtained in the other conditions. We found that the power law accounted well for the results irrespective of filtering (r2>0.85 over the tested range of frequency cutoffs). The value of β-exponent varied with frequency cutoff, but only to a limited extent (median=0.77, interquartile-range=0.06). We also computed the least-squares orthogonal regression of log A(t) versus Log C(t) forthe tail of the larvae. Across all animals and conditions (overshoot, approach, dispersal),the median variance accounted for by the power law (r2) was 0.89 (interquartile-range=0.06, n=123), and the median value of the power-exponent was 0.74 (interquartile-range=0.09).

## Results

Locomotion of individual larvae on a flat agar surface was tracked at a high spatio-temporal resolution (**Materials and Methods**). To induce animals to naturally ‘draw’ different types of trajectories, we tested different sensory environments [8]. In the overshoot condition during chemotaxis close to an odor source, the larval centroid traced complex trajectories resembling human scribbles (**Figure 1A**). Trajectories were not associated with a constant progression speed or any simple kinematic pattern. A log-log plot of speed versus curvature revealed a power law as a straight line whose slope corresponds to the power-exponent (**Figure 1B**). Both the instantaneous angular velocity and local path curvature were widely modulated, yet they co-varied throughout (**Figure 1C**). Similar results were obtained for all individual larvae in this condition. Next, we tested larvae subjected to other sensory conditions, resulting in different exploratory strategies and movement trajectories. In the approach condition, individuals reached an odor source at the opposite side of the arena via progressively more curved paths(**Figure 1D**). In the dispersal condition, larvae searching for food in the absence of olfactory cues moved in arbitrary directions tracing highly variable paths (**Figure 1G**). Overall, the power law did not depend on the type of exploratory movements: overshooting, approaching and dispersing larvae all complied with the power law (compare **Figures 1B,E,H**, and **Figure I**).

**Fig. 1.**
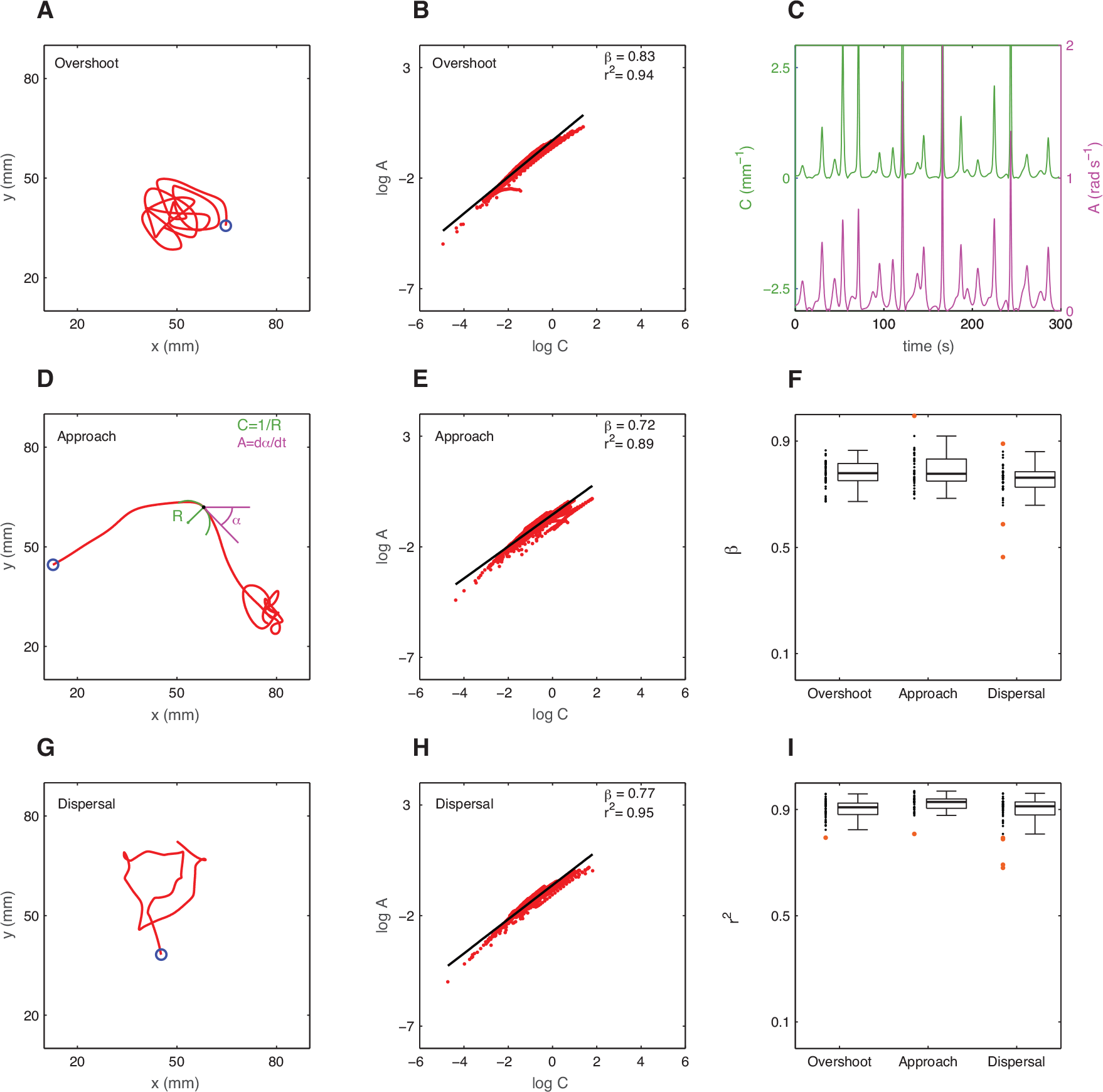
Speed-curvature relation in *Drosophila* larvae tracing different trajectories. (**A**)Trajectory of the centroid position of one representative larva in the overshoot condition (blue circle indicates starting position). (**B**) Scatter-plot of instantaneous angular velocityand local curvature on a loglog scale. All data points (red dots, n=2100) sampled at equal time intervals along the same trajectory as in panel A were included. The data were fitted by the power function A(t)=kC(t)^β^ (black line) with β-exponent and r2 variance accounted for as indicated in the inset. (**C**) Temporal evolution of the local path curvature (green) and instantaneous angular velocity (maggenta) for the same data as in A-B. (**D**) Centroid trajectory of a larva in the approach condition. Key movement variables are identified at an arbitrary point along the trajectory: C is the curvature of the osculating circle of radius R, α is the phase angle of the tangent, and the angular velocity A is the time-derivative of α. (**E**) Loglog plot of angular velocity versus curvature for the same trajectory as in D. (**F**) Summary boxplot statistics for β-exponent of individual animals in the 3 different groups: overshoot(n=42), approach (n=40), and dispersal (n=41). Outliers are orange dots. (**G**) Centroid trajectory of a larva in the dispersal condition. (**H**) Log-log plot of angular velocity versus curvature for the same trajectory as in G.(**I**) Summary boxplot statistics for r2 in the 3 groups.

The median value of the power-exponent was 0.78 (interquartile-range=0.06, n=42), 0.78 (interquartile-range=0.08, n=40), and 0.76 (interquartile-range=0.06, n=41) for the overshoot, approach, and dispersal conditions, respectively (**Figure 1F**). The distribution of the power-exponentsdid not differ significantly between the 3 groups (Kruskal-Wallis H(2,123)=5.29, p=0.071; multiple comparisons p>0.05). Similar results were observed for trajectories traced by the animal’s hindmost part rather than the centroid (**Materials and Methods**). Moreover, the results were not affected substantially by using different frequency cutoffs in filtering the position data (**Figure S1**). A few individuals did not comply with the law (especially in the dispersal condition, see outliers in **Figures 1H,I**), corroborating that it is not an obligatory outcome of our analyses. For the sake of comparison, the exponent for human hand-drawing is generally close to 0.66 (so called2/3-power-law [1]), but it becomes 0.73 when drawing in water [12], the latter value being close to the present values in larvae. Therefore, not only do we find in the larvae the geometric-kinematic constraint dictated by the power law, but also hints of dynamic constraints in the exponent [2,5,12], as recently found in humans, where the specific value of the exponent appears to depend on the viscosity of the medium (air or water for hand-drawing,agar support and thin liquid coat for larvae).

## Discussion

In *Drosophila* larvae, multiple central pattern generators in the abdominal and thoracic segments of the nervous system generate peristaltic waves of muscle contractions along the body axis that enable crawling [6]. The degree of symmetry and synchrony of neural activity on each side of the nervous system might control the instantaneous direction of movement, straighter trajectories resulting from more symmetrical contractions in amplitude and timing [13],while frequency might determine movement speed. It is then possible that the velocity-curvature power law emerges from these patterns of neural activity transformed in oscillatory body motion. However, suprasegmental nervous structures as well as sensory feedback also contribute to the net motor output [6]. Future studies might use appropriate mutants to genetically disrupt [13] the power law to elucidate the relative role of neural structures, sensory feedback, and body mechanics. The possibility of genetic manipulations, electron microscopy reconstructions, electrophysiology and imaging settle the larva as a privileged system to dissect the mechanistic implementation of the law.

Without overstressing similarities for translational purposes, nor dismissing superficial differences, here we have reported the discovery that a fundamental law of human control is at work in the humble maggot. The power law for voluntary movements in human and non-human primates may well have different origins [4] from those in crawling fruit-fly larvae. Yet, it is remarkable that the law is compatible with relatively much simpler nervous systems, and that it holds for movements differing in speed by several orders of magnitude, as those generated by humans and maggots. Scaling laws such as the velocity-curvature power law might exist because natural selection favors processes that remain behaviorally efficient across wide ranges of size and structure in different contexts in distantly related species [3,7]. The present work is an example of the search for shared principles of animal behavior across phyla.

## Acknowledgments

We thank Asif Ghazanfar, André Brown and Santiago Canals for valuable feedback.

## Author contributions

All authors contributed to the design of the study, analysis of data, interpretation of results, and the final version of the manuscript.

---

**Fig. S1.**
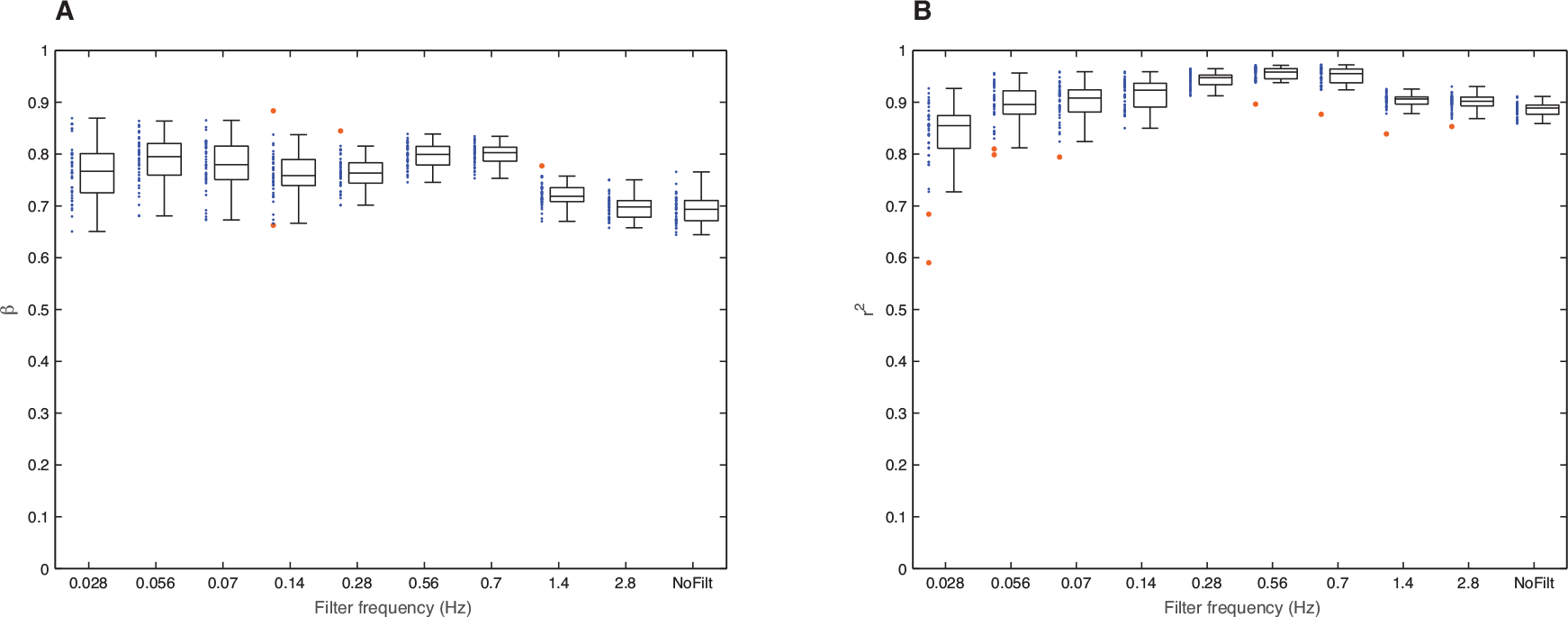
Effect of low-pass filtering on the power law. The power function A(t)=kC(t)^β^ was fitted to the results of the larvae in the overshoot condition, after filtering the raw x,y position samples of the centroid at the frequency cutoff indicated on the abscissae. *Nofilt* denotes no filter whatsoever. Summary boxplot statistics are plotted for the β-exponent (**A**) and r2 (**B**). Outliers are orange dots.

